# Norepinephrine, olfactory bulb and memory stability

**DOI:** 10.1101/2020.06.17.153502

**Authors:** Christiane Linster, Maellie Midroit, Jeremy Forest, Yohann Thenaisie, Christina Cho, Marion Richard, Anne Didier, Nathalie Mandairon

## Abstract

Memory stability is essential for animal survival when environment and behavioral state change over short or long time spans. The stability of a memory can be expressed by its duration, its perseverance when conditions change as well as its specificity to the learned stimulus. Using optogenetic and pharmacological manipulations, we show that stability of an odor-reward association can be modulated by noradrenergic inputs to the first olfactory network, the olfactory bulb. We show that while manipulations of noradrenaline during an odor-reward acquisition have no acute effects, they impact learning flexibility as well as the duration and the specificity of the memory. We use a computational approach to propose a proof of concept model showing that a single, simple network effect of noradrenaline on olfactory bulb dynamics can underlie these seemingly different behavioral effects. Our results show how acute changes in network dynamics can have long term effects that extend beyond the network that was manipulated.

## Introduction

Stability to maintain skills and stimulus associations while adjusting to new circumstances is fundamental for animal behavior. Neural systems need to both be flexible to adapt rapidly to new information and stable to maintain learned behaviors to ensure survival. Neuromodulatory systems in particular are thought to have evolved to convey flexibility to neural systems by allowing them to process stimuli in very different manners depending on behavioral demands.

For example, acetylcholine may sharpen stimulus evoked oscillations in neural networks to enhance attention to a stimulus on a momentary basis (Sarter et al., 2006, Parikh and Sarter, 2008), dopamine may increase plasticity when a stimulus is unexpected and in need to be reinforced (Gentry et al., 2019), or norepinenphrine (NE) may increase signal to noise ratio in specific networks during moments of stress to enhance recognition and processing of a stimulus (McBurney-Lin et al., 2019). NE has long been associated with olfactory learning (Devore and Linster, 2012, Linster and Escanilla, 2019), and been shown to strongly modulate the processing of olfactory stimuli as early as in the olfactory bulb (OB). NE inputs to the OB change processing of low amplitude odors, increase associative learning and decrease signal to noise ratio (Doucette et al., 2007, Escanilla et al., 2010, Devore and Linster, 2012, Escanilla et al., 2012, Linster and Escanilla, 2019). We here show that in addition to these acute effects, temporary manipulations of bulbar NE modulate long term stability of olfactory memories.

## Results and Discussion

We first show that memory stability depends on bulbar NE by optogenetically decreasing NE fiber activity while testing mice on a reversal task and a long-term memory task. We then use a pharmacological approach to narrow down NE receptor types underlying our observations. Last, we show that memory stability as expressed by the specificity of this memory is modulated by bulbar NE and use a computational approach to propose a single mechanism that could underlie all of these observations.

### (1) NE modulation in the olfactory bulb increases odor memory stability expressed as perseverance in response to change

This experiment tested how stable a memory acquired after a training period of10 trials is by using a reversal training paradigm. To inhibit NE transmission specifically in the OB, hSyn-eNpHR3.0-EYFP lentivirus was injected bilaterally into the Locus Coeruleus and bilateral optical fibers were implanted into the OBs(NpHR mice). Control mice were injected with hSyn-EYFP lentivirus and similarly implanted with optical fibers (Ctrl mice). The inhibition of NA fibers in the OB was performed by bilateral continuous light stimulation (561 nm, 10–15 mW) automatically triggered by the entry of the mouse’s nose 5 cm around the odorized holes (VideoTrack, Viewpoint). Expression of light sensitive chloride pumps and delivery of light to axonal projection targets has been successfully used to inhibit activity at presynaptic terminals (Mahn et al., 2016; Spellman et al., 2015; Stuberet al., 2011; Tye et al., 2011; Raimondo et al.,2012; Restrepo 2018). NpHR and Ctrl mice were trained under light stimulation in a 10-trial simultaneous go-no-go task (Escanilla et al., 2008, Chaudhury et al., 2009, Moreno et al., 2012, Mandairon et al., 2018) in which they had to associate odorant 1 with a reward while odorant 2 was not reinforced (Odorset 6; Supplementary Table 1). These 10 trials were immediately followed by 15 trials of reversal learning with no light stimulation in which odorant 1 was not reinforced and odorant 2 was rewarded (Figure 1A). The latency to find the rewarded odorant was recorded as a measure for how well the odor-reward association was learned. Both experimental groups rapidly acquired the odor-reward association as evidenced by significant decreases in delay to find the rewarded odor (see Table 1 for statistical results, Figure 1A). During reversal learning, control mice showed a high degree of perseverance to the previously learned association even after 15 trials, whereas NpHR mice quickly learned the new odor-reward association (trials 6-10; Figure 1A and Table 1). The efficiency of viral transduction in terminal fibers was assessed by calculating in the OB the percentage of double labeled EYFP-positive and NE-positive fibers in the OB (labeled by immunohistochemistry of NE-transporter; NET) among NET-positive fibers on all animals of both experimental groups. (Supplementary Figure 1).

**Table 1:** Summary of statistical analyses

**Reversal training experiment**
| Repeated measures ANOVA | Acquisition* (Wilk's Lambda)<br>Trials 1-5 versus 6-10 | Reversal ** (Wilk's Lambda)<br>Trials 6-10 versus 21-25 |
| --- | --- | --- |
| $F_{\text{trialblock}}(4,3) = 15.958; p < 0.023$<br>$F_{\text{trialblock} \times \text{group}}(4,3) = 5.835; p = 0.049$ | Control: $p = 0.002$<br>NpHR: $p = 0.042$ | Control: $p = 0.268$<br>NpHR: $p = 0.034$ |

\* Significant decrease of delay between trials 1-5 and 6-10 indicates learning
\*\* Significant increase between end of acquisition (trials 6-10) and end of reversal (21-25) indicates lack of reversal learning

**Long-term memory experiment**
| Repeated measures ANOVA | Acquisition* (Wilk's Lambda)<br>Trials 1-5 vs 16-20 | 2 hour Recall ** (Wilk's Lambda)<br>Trials 1-5 vs 2hrs | 24 hour Recall *** (Wilk's Lambda)<br>Trials 1-5 vs 24hrs |
| --- | --- | --- | --- |
| Acquisition (trials 1-20)<br>$F_{\text{trialblock}}(3, 20) = 3.614; p = 0.186$<br>$F_{\text{trialblock,group}}(3, 20) = 0.915; p = 0.081$<br>Recall (2 and 24 hour trials)<br>$F_{\text{trialblock}}(1,19) = 15.345; p = 0.001$<br>$(F_{\text{trialblock,group}}(1, 19) = 10.121; p = 0.048$ | Control: $p = 0.015$<br>NpHR: $p = 0.004$ | Control: $p = 0.041$<br>NpHR: $p = 0.010$ | Control: $p = 0.023$<br>NpHR: $p = 0.754$ |
| | | | 2 hour vs 24 hour\$<br>Control: $p = 0.649$<br>NpHR: $p < 0.001$ |

\* Significant decrease of delay between trials 1-5 and 6-10 indicates learning
\*\* Significant decrease between trials 1-5 and 2 hrs indicate recall
\*\*\* Significant decrease between trials 1-5 and 24 hours indicate recall

**Pharmacology long-term memory experiment**
| Repeated measures ANOVA | Acquisition* (Wilk's Lambda)<br>Trials 1-5 vs 16-20 | 24 hour Recall *** (Wilk's Lambda)<br>Trials 1-5 vs 24hrs |
| --- | --- | --- |
| $F_{\text{trialblock}}(4, 32) = 13.354; p < 0.001;$<br>$F_{\text{trialblock} \times \text{drug}}(16, 98.399) = 1.753; p = 0.026$ | Saline: $p = 0.042$<br>Alpha1: $p = 0.035$<br>Alpha2: $p = 0.021$<br>Beta: $p = 0.029$<br>All: $p = 0.001$ | Saline: $p = 0.023$<br>Alpha1: $p = 0.466$<br>Alpha2: $p = 0.019$<br>Beta: $p = 0.021$<br>All: $p = 0.723$ |

\* Significant decrease of delay between trials 1-5 and 6-10 indicates learning
\*\* Significant decrease between trials 1-5 and 24 hours indicate recall

**Memory specificity experiment**
| Repeated measures ANOVA | Perceptually similar odor *<br>C versus C+1 | Perceptually less similar odor *<br>C versus C+2 | Unrelated odor *<br>C versus X |
| --- | --- | --- | --- |
| $F_{\text{odor}}(3, 28) = 13.023; p < 0.0001;$<br>$F_{\text{odor} \times \text{drug} \times \text{trials}}(6, 58) = 1.2222; p = 0.310$ | Saline:<br>4 trials: $p = 0.299$<br>8 trials: $p = 0.039$<br>12 trials: 0.008<br>Phentolamine:<br>4 trials: $p = 0.835$<br>8 trials: $p = 0.357$<br>12 trials: $p = 0.039$ | Saline:<br>4 trials: $p = 0.029$<br>8 trials: $p = 0.003$<br>12 trials: 0.001<br>Phentolamine:<br>4 trials: $p = 0.502$<br>8 trials: $p = 0.032$<br>12 trials: $p = 0.040$ | Saline:<br>4 trials: $p = 0.045$<br>8 trials: $p = 0.011$<br>12 trials: 0.016<br>Phentolamine:<br>4 trials: $p = 0.005$<br>8 trials: $p = 0.004$<br>12 trials: $p = 0.04$ |

\* Significant decrease of digging during unrewarded trials indicates discrimination between C and novel odor (with Wilk's lambda)

**Figure 1.**
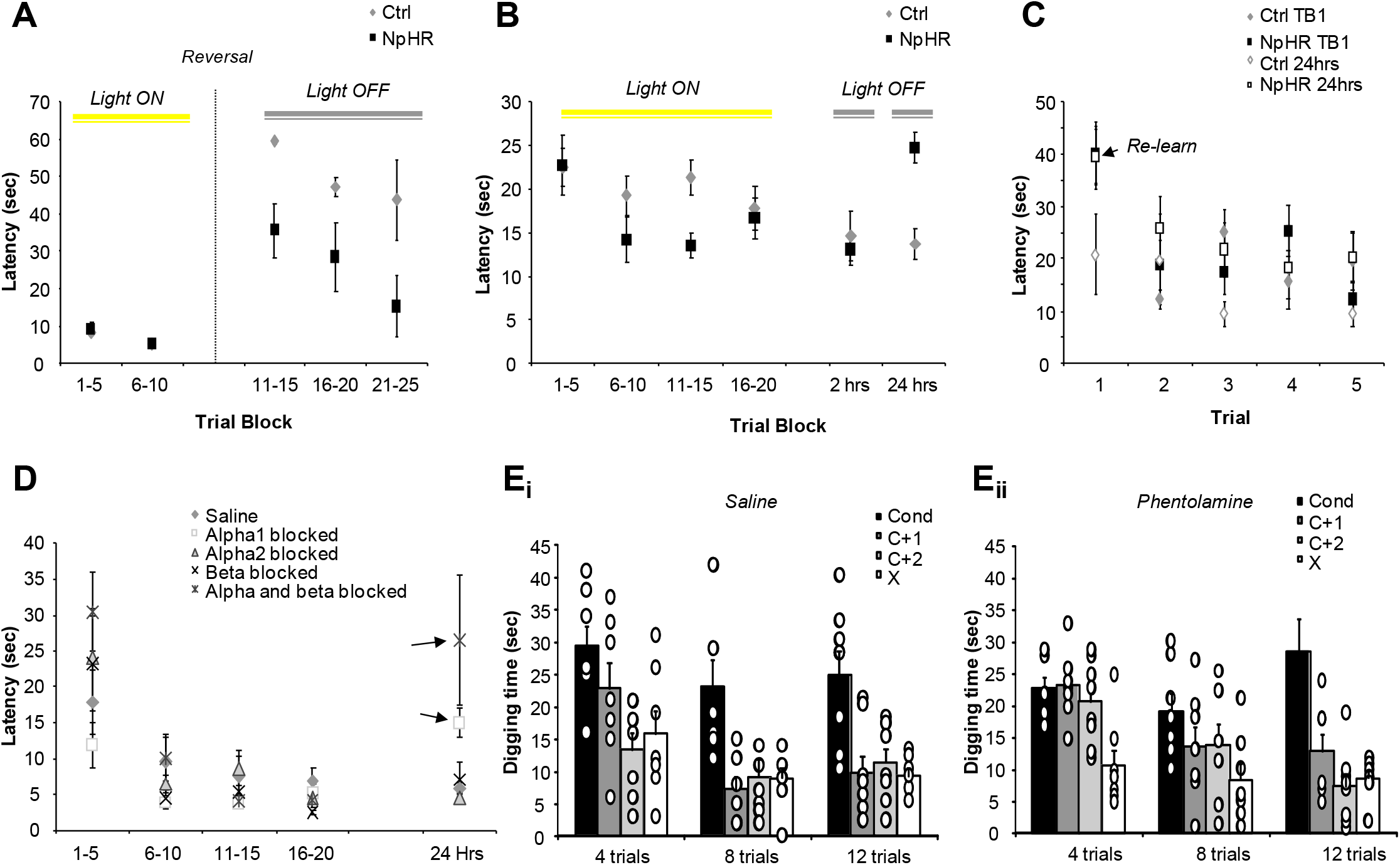
Decreased NE inputs to the OB renders odor memories less stable. **A**. Reversal experiment. The graph shows the average delay to find the rewarded odor as a function of trial block. Odor-reward contingency was reversed after 2 trial blocks (reversal). Optogenetic blockade of NE inputs was performed only during the initial 10 trials (trial blocks 1-5 and 6-10, Light ON), not during reversal training. Control mice with NE modulation not affected by light stimulation did not learn the new odor-reward association whereas NpHR mice (NE modulation decreased by optogenetic stimulation) acquired the new association after 20 trials. **B**. Duration of odor memory is shortened when bulbar NE is decreased. The graph shows the average delay to find the rewarded odor as a function of trial block for both experimental groups. Optogenetic decrease of NE inputs to the OB (Light ON) was performed during 20 acquisition trials (trial blocks 1-5, 6-10, 11-15, 16-20) but not during recall trials 2 hours and 24 hours after the end of acquisition (Light OFF). Note that in NpHR animals the delay to find the rewarded odor was increased to the level during initial acquisition at the 24 hour test block. **C**. Experimental animals re-learn the task at the 24 hour delay block. The graph shows average delays to find the rewarded odor during the first 5 acquisition trials (Ctrl 1-5 and NpHR 1-5) and during the five trials at the 24 hour recall test (Ctrl 24hrs and NpHR 24 hrs). Note that NpHR mice perform similarly during these two blocks showing that they re-learn rather than recall the odor-reward association. **D**. NE effects on memory duration are mediated by OB alpha1 receptors. The graph shows acquisition (trials 1-20) of the odor-reward association during local pharmacological blockade of NE receptors in the OB (no blockade, alpha1, alpha2, beta or all receptors blocked) and 24 recall with no receptor blockade. Note that mice with alpha1 or all receptors blocked showed significantly longer delays for recall after 24 hours than saline controls, showing that NE effects on long term memory are mediated at least partially by alpha1 receptor activation. **E**. Effect of alpha NE receptor blockade on acquisition and memory specificity. The graphs show the time mice dig in an odor (conditioned or novel odors of decreasing similarity) during unrewarded test trials after 4, 8 or 12 acquisition trials. Note that saline infused control mice (**Ei**) were able to discriminate less similar odors (C+2, X) from the conditioned odor after just 4, and all novel odors after 8 conditioning trials, whereas mice with NE receptors blocked (**Eii**) needed 12 conditioning trials to discriminate all novel odors.

<u>These results show that the stability of the acquired memory is weaker for NpHR as compared to Ctrl animals; while effects of inhibiting bulbar NE are not evident during the initial acquisition of an odor-reward association, they manifest during reversal training even though NE is not modulated at that time.</u>

### (2) NE modulation in the olfactory bulb increases odor memory stability expressed as an increase in memory duration

To test how the presence of NE during learning affects the duration of the odor memory, we trained the same mice for 20 acquisition trials on the same task (Odorsets 3&4, Supplementary Table 1). We tested their recall ability 2 hrs and 24 hrs later (Figure 1B). NE transmission in the OB was light inhibited during the acquisition but not recall trials. We found that both experimental groups decreased their latency to find the rewarded odorant over the course of 20 trials with no difference between the groups (Figure 1B and Table 1). Both groups were equally quick at finding the rewarded odorant 2 hours after the initial training; however, 24 hours after training, latencies to find the reward were significantly increased in NpHR mice (Figure 1B and Table 1). We compared performance during the first five acquisition trials and the five trials at 24 hours: while control mice had significantly lower delays during the 24 hour trials as compared to the initial acquisition (F(1, 8) = 7.9; p = 0.023), NpHR mice have similar delays in both cases (F(1, 11) = 3.037; p = 0.109) showing they relearned the task (Figure 1C). A control experiment showed that NE inhibition during 2 hour or 24 hour recall did not affect the delays to find the correct odorant (Supplementary Figure 2); this shows that release of NE blockade did not have an effect on odor perception or task performance per se.

<u>These results show that while inhibition of bulbar NE release does not noticeably slow down acquisition of an odor reward association, it does impair the duration of memory, confirming our hypothesis that the presence of NE during acquisition renders a memory more stable.</u>

We used a pharmacological approach to identify which NE receptors in the OB are mediating this effect. Mice (n=12) were implanted with billateral cannulae in the OB for intracerebral drug infusions and tested using the same paradigm. In each training session mice were trained for 20 trials with either alpha1, alpha2, beta, all or no NE receptors blocked (Mandairon et al., 2008) and tested 24 hours later with a single 5 trial block and no drug infusions (Figure 1D). Results showed that all mice acquired the task similarly, but that mice with alpha 1 receptors blocked during acquisition showed significantly longer delays to find the rewarded odor 24 hours after training (Table 1 and Figure 1D). <u>Alpha 1 receptor blockade locally in the OB had similar effects than light-inhibition of NE fiber activity locally in the OB, showing that NE effects on long term memory are mediated at least partially by alpha 1 receptor activation.</u>

### (3) NE modulation in the olfactory bulb and amount of training increase odor memory stability as expressed in specificity for the learned odor

Experiment 3 tested to what degree the amount of training as well as activation of NE receptors determine the specificity of an odor-reward association. To test memory specificity, we used a generalization talk in which mice learn to associate an odor with reward and are later tested on novel odors to assess how specific the formed memory is (Cleland et al., 2002, Cleland and Narla, 2003, Mandairon et al., 2006, Chaudhury et al., 2009, Cleland et al., 2009). Mice (n=9) were implanted with bilateral cannulae and trained to associate an odorant with a reward during 4, 8 or 12 trials. After completion of training trials, mice were tested in 4 consecutive counterbalanced unrewarded trials with the conditioned odor (C), two chemically and perceptually related novel odors (C+1, C+2) and one unrelated novel odor (X). How long mice search for the reward in a novel odor is a measure for how much they confuse the novel odor with the conditioned odor and hence assesses memory specificity (Linster and Hasselmo, 1999). Mice were trained after infusion of saline or the non specific alpha receptor blocker phentalomine into both OBs.

Saline treated control mice showed higher specificity for the conditioned odor when trained longer: the perceptually most similar odor (C+1) was discriminated after 8 or 12 training trials only, while the unrelated odor (X) was discriminated after as few as 4 trials. In contrast, mice with NE receptors blocked needed 12 training trials to discriminate the two perceptually similar odors (C+1 and C+2) strongly suggesting that the presence of NE during learning enhances memory specificity (Figure 2 and Table 1).

**Figure 2.**
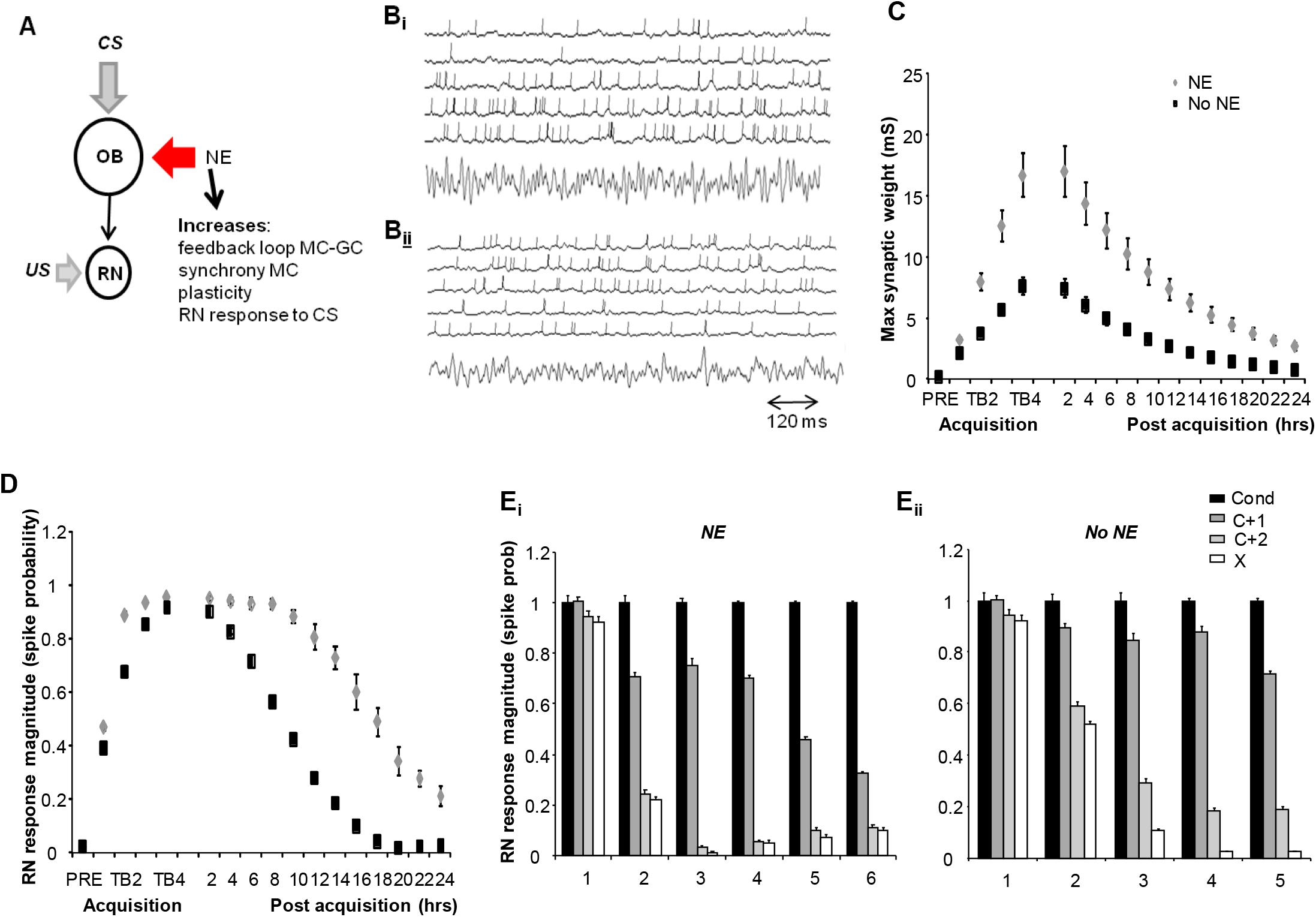
Computational modeling of bulbar NE effects on learning. **A**. Schematic illustration of the model structure. Olfactory stimuli are sent to OB network via activation of simulated olfactory sensory neurons. The OB network also receives NE modulatory inputs (see methods). MC cells make plastic excitatory synapses with a response neuron (RN) which also receives input when a reward (US) is paired with an odor. **B**. Neural activity and field potentials in the model with (**B_i_**) and without (**B_ii_**) NE modulation. The traces show action potentials and voltage fluctuations of representative OB neurons in the model, the lower trace shows the simulated field potential. Note that synchrony among spikes and field potential dynamics are decreased when NE modulation is impaired in the model. **C**. Changes in the maximal synaptic weights between MCs and RN. The graph shows the strength of synaptic weights (in mS) as a function of trial block during acquisition and hours elapsed during forgetting (post acquisition in hours) for simulations with and without NE modulation. Note that when NE is omitted during acquisition, synaptic weights increase less and decreases to initial values by 18 hours, leading to non-recall at the 24 hour time point (compare to D). **D**. Response of RN (spike probability) to OB stimulation with the conditioned odor during acquisition and forgetting. RN response magnitude was measured after every trial block during acquisition and after every simulated hour during post acquisition (without US). Note that when NE modulation is omitted during acquisition, RN responses increase to the same asymptotic value, however, responses decrease more rapidly post acquisition and by 18 hours post acquisition RN responses are as low as baseline. **E**. RN response specificity as a function of amount if training with (**E_i_**) and without (**E_ii_**) NE. The graphs show the ratio of RN response magnitude to novel odors N1-N3) as compared to the conditioned odor (C) before training (Pre) and after 1, 2, 3 or 4 trial blocks. Note that specificity, indicated as responses to novel odors being lower than response to conditioned odor always increases with trial blocks, but this effect is more pronounced when NE modulation during training is simulated. RN responses are tested after each trial block in the absence of reward inputs.

<u>Results from Experiment 3 show that specificity for an odor memory increases when NE is unimpaired in the OB. Both the number of trials and the presence of NE modulated the stability of the formed memory, as expressed by its specificity to the learned odor.</u>

In a separate experiment, we tested the idea that amount of training would increase stability also with respect to memory duration (Supplementary Figure 3). Mice were trained for 5, 10, 15 or 20 trials and tested 2 hours or 24 hours later. We found a strong correlation between memory duration and the amount of training (Supplemental Figure 3).

### (4) Olfactory bulb NE enhances synchrony and thereby learning, leading to increased memory stability, duration and specificity

We present a computational model of NE modulation in the OB and odor-reward association learning as proof of concept that the diverse long term effects of bulbar NE we observed can be mediated by its effects on bulbar dynamics. In a nutshell, higher synchrony at the output of the OB results in enhanced plasticity, mediating stronger, longer lasting and more specific odor reward associations.

Our simplified model (Figure 3A) implements NE modulation as measured in brain slice experiments (Nai et al., 2009, Nai et al., 2010, Linster et al., 2011, Linster, 2019) and in vivo (Manella et al., 2017) and determined computationally (Escanilla et al., 2010, Linster et al., 2011, de Almeida et al., 2015). Briefly, NE inputs to the OB model enhanced mitral cell (MC) excitability and granule cell (GC) spontaneous activity levels, resulting in stronger MC-GC interactions and more pronounced MC spike synchronization and stronger, higher contrast responses to simulated odors (Figure 3B; quantified in Escanilla et al. 2010). This results in stronger learning of the odor-reward association, as evidenced by a larger increase in synaptic weights (Figure 3C) and response to the conditioned stimulus (CS, odor) by the response neuron (RN; Figure 3D). Interestingly, with the parameters we chose here, the response to the CS, the odor, assessed as the level of activation of the RN, is not much affected by the presence of NE during acquisition (Figure 3D), but is evident in post testing of response levels post acquisition and during response specificity testing (Figure 3E), similar to what we saw behaviorally.

In the simulations, NE during acquisition of the odor-reward association has multiple effects. First, stronger synaptic weights in response to the conditioned odorant when acquired with NE (Figure 3C) implement a “stronger” memory not as easily reversed (as shown in Experiment 1). Second, NE in the OB during acquisition increases memory duration simply because of increased synaptic weights after 4 trial blocks of learning (Figure 3C): RN responses to the conditioned odor decrease to baseline before the 24 hour recall test when NE was omitted (as shown in Experiment 2). NE in the OB during acquisition increased specificity of the learned RN response as a function of training trials: when NE modulation was simulated during acquisition, responses of the RN to novel odorants rapidly decreased as learning increased, with even the most overlapping stimulus significantly discriminated after 2 trial blocks (comparable to behavioral data; Figure 3Ei). In contrast, when acquisition in the model OB was simulated without NE, discrimination of the most similar odor required 4 trials blocks (Figure 3Eii). This effect results from synaptic weight distributions becoming more specific to the odor as learning proceeds (Supplementary Figure 3).

Our simulations show how modulation of bulbar dynamics by NE can have multiple long term effects on odor-reward association. We created a very simplified model meant to provide proof of concept rather than a detailed model of how odor reward associations would happen.

## Conclusion

Our experimental results show that stability of an olfactory memory, measured by its perseverance, duration and specificity, is enhanced when NE release and activity is undisturbed in the OB. A computational model incorporating known cellular and network effects of NE in the OB suggests that NE modulation of OB dynamics are mediating these effects. Interestingly we do not observe acute effects of lacking NE activity during learning, but see significant effects of this temporary manipulation long after the manipulation has stopped, which can underlie observations with respect to NE and stress in the OB (Manella et al., 2013) and NE and integration of adult born neurons into the network (Moreno et al., 2012).

## Material and Methods

### Optogenetic experiments

#### Animals

12 adult male C57Bl6/J mice (Charles River Laboratories, L’Arbresle, France) aged 2 months at the beginning of the experiments were used for this experiment. Mice were housed in standard laboratory cages and were kept on a 12 hr light/dark cycle (at a constant temperature of 22°C) with food and water ad libitum except during behavioral tests during which they were food deprived (∼20% reduction of daily consummation, leading to a 10% reduction in body weight). Mice were housed by group of 5, and individually after surgery. All experimental procedures were validated by Lyon 1 and the French Ethical Committee (protocol n° DR2013-48).

#### Odorants

Seven pair of odorants were used in this study. Odorants diluted in mineral oil to achieve an approximate gas phase partial pressure of 10 Pa (Cleland et al., 2002, Kermen et al., 2011); Supplementary Table 1)

#### Viral vector injection and optical fiber implantation

Prior to surgery, mice were anesthetized with a cocktail injection of 50 mg⁄kg ketamine and 7.5 mg⁄kg xylazine (i.p.) and secured in a stereotaxic instrument (Narishige Scientific Instruments, Tokyo, Japan). 300 nl of hSyn-eNpHR3.0-EYFP lentivirus (9,22 × 106 IU/ml, expressing halorhodopsin and the yellow fluorescent protein; n=7) and 300nl of control hSyn-EYFP lentivirus (1,1 × 106 IU/ml, expressing only EYFP; n=5) were injected bilaterally into the Locus Coeruleus at the following coordinates with respect to the bregma: AP, + −5.4 mm; ML, ± 0.9; DV, - 4 mm at a rate of 150 nl/min. In all mice, bilateral optical fibers (200-nm core diameter, 0.22 N.A.; Doric Lenses) were implanted into the OBs (from bregma : AP, + +4.6 mm; ML, ± 0.75 mm; DV, −2 mm). Behavioral experiments were performed 8 weeks after surgery. The pLenti-hSyn-eNpHR 3.0-EYFP was a gift from Karl Deisseroth (Gradinaru et al., 2010) and obtained through Addgene (plasmid #26775). Elaboration of the control pLenti-hSyn-EYFP lentivirus has been previously described (Kermen et al., 2016).

#### Behavioral procedure

##### Apparatus

Behavioral training took place on a computer assisted two-holes board apparatus described previously(Mandairon et al., 2009). The hole-board is equipped with capacitive sensors that monitor the events of nose-poking (visits) in the holes. The holes were odorized by placing a cotton swab impregnated with 60 μL of 10 Pa odorant under bedding in a small dish placed into the hole. A food reward was buried into the bedding of one of the holes, with the location of the odor-reward randomly determined for each trial.

##### Shaping

The mice were first trained to retrieve a reward (a small bit of sweetened cereal; Kellogg’s, Battle Creek, MI, USA) by digging through the bedding. The mouse was put in the start area of the two hole-board apparatus and allowed to dig for 1 min. During the first few trials, the reward was placed on top of the bedding in one of the holes. After the mice successfully retrieved the reward several times, it was successively buried deeper and deeper in the bedding. Shaping was considered to be complete when a mouse could successfully retrieve a reward buried deep in the bedding for a least 16 out of 20 trials. Odor set 1 was used for shaping.

##### Learning and memory tests

Each session consisted of one minute trials during which the mouse was allowed to retrieve the food reward from thehole. If a mouse failed to find the reward after 60 seconds, the trial was ended and the mouse replaced on the starting position behind a cover while the next trial is set up. The inhibition of NAfibers in the OB was performed by bilateral continuous light stimulation (crystal laser, 561 nm, 10–15 mW) automatically triggered by the entry of the mouse’s nose within a 5 cm zone around the odorized hole (light-triggering zone; VideoTrack, Viewpoint) and stopped automatically when the mouse’s nose exited the zone.

##### Reversal test

Mice were first trained for 10 trials of 1 min on an odor-reward association immediately followed by 15 trials with reversed odor-reward contingency. Optical stimulation was used for the initial 10 acquisition trials only. Mice were tested using odor set 6 with O1 associated with the reward in the first 10 trials and O2 associated with the reward for the last 10 trials. Each mouse was tested once.

##### Long term memory test

Mice were trained with optical stimulation during the 20 acquisition trials and tested without optical stimulation for 5 trials 2 hours and 24 hours later. Each mouse was tested twice in this experiment, once with odorset 3 and once with odorset 4, with O1 for each odorset associated with reward. A control experiment tested the role of NE inputs to the OB for recall: mice were trained during 20 trials without optical fiber stimulation and tested at 2 and 24 hours with optical stimulation. Each mouse was tested once in this experiment, using odorset 5, with O1 associated with reward.

##### Statistical analysis

All statistical analyses were performed using SPSS. Analysis was performed on the latency (delay) to dig in the rewarded dish as dependent variable. The latency to dig in the rewarded dish is a good indicator for the strength of the acquired memory, with short latencies signaling a strong memory and fast decision making and longer latencies signaling slower decision making and weaker learning (Mandairon et al., 2018). Repeated measures ANOVA were used to assess how blockade of NE projections modulates acquisition or recall. In each case, experimental group (control or NpHR) was used as between subjects factor and trial block or trial number as within subjects factor (alpha = 0.05). Pairwise comparisons between trial blocks to assess acquisition and recall were performed using Wilk’s Lambda with alpha=0.05.

#### Cellular analysis

Mice were sacrificed using pentobarbital (0.2 ml/30 g) and intracardiac perfusion of 50 ml of fixative (PFA 4%, pH = 7.4). The brains were removed, post-fixed overnight, cryoprotected in sucrose (20%), frozen rapidly, and then stored at −20°C before sectioning with a cryostat.

Immunohistochemistries of NET (NE-Transporter to label NE fibers) and EYFP (to label transduced fibers) were performed in 4-6 section (40 μm thick) of the OB distributed along its antero posterior axis, using anti-NorEpinephrin Transporter (mouse, Mab technologies; 1/1000) and anti GFP (chicken, Anaspec/Tebu; 1/1000). Appropriate secondary antibodies were used (goat anti-mouse Alexa 546 Vector; 1/250 and goat anti-chicken Alexa 488, Molecular Probes, 1/250). Sections were then cover-slipped in Vectashield (Vector laboratories).

All fluorescent analyses were done blind with regards to the identity of the animal. Images were taken in the granule cell layer of the OB with a Zeiss microscope equipped with an apotome, using 40 oil-immersion objective. Z-stacks were acquired with 0. 2 μm interval between images. 8-13 pictures per animal were analyzed. Length of NET- and NET/GFP-positive fibers were analyzed with 3D viewer of ImageJ. A percentage of NET/GFP-positive fibers among NET-positive fibers was assessed. No animal was excluded from the analysis. After testing the normality and variance of the data, bilateral unpaired parametric t-test was performed.

### Pharmacological experiments

#### Animals

12 (Experiment 2) and 9 (Experiment 3) adult male C57Bl6/J mice (Charles River Laboratories) aged 2 months at the beginning of the experiments were used for pharmacological experiments. Mice were housed in standard laboratory cages and were kept on a 12 hr light/dark cycle with food and water ad libitum except during behavioural test where they were food deprived to no less than 85% of their free feed weight. All experimental procedures were conducted under a protocol approved by the Cornell University IACUC.

*Odorants* were those listed in Supplementary Table 1. Before each behavioral session, 60 uL odor was loaded onto 5mL sand and then covered with additional 5mL of sand.

#### Cannulation surgery

After behavioral shaping and prior to experiments, mice underwent surgery to implant bilateral cannulae in the OB for drug delivery according to established methods. Mice were anesthetized with gas anesthesia (isoflurane, 2-4%), injected intraperitoneally with 0.05mg/ kg atropine, and guide cannulae (22-gauge, Plastics One) were inserted 5mm anterior and 1.5 mm ventral from Bregma and affixed to the skull with dental cement(Guerin et al., 2008, Tong et al., 2018). After surgery mice were given pain killers and saline injections and allowed to recover for 7 days.

#### Drug infusions

20 minutes before behavioral testing, mice were bilaterally infused with NE antagonists or saline. 2 uL of solution was infused at 1 uL/minute and the infusion needle was left in place for 5 minutes after infusion. For long term memory testing, mice were infused with saline, the alpha 1 blocker prazosin (hydochloride, 1mM), the alpha 2 blocker yohimbine (hydochloride, 2mM), the beta antagonist alprenolol (hydochloride, 12mM) or a cocktail of all three, all purchased from sigma and diluted in 0.9% saline. For specificity testing, mice were infused with the non-specific alpha antagonist phentolamine (12mM) or saline, with dosages determined from our previous behavioural experiments (Mandairon et al., 2008, Escanilla et al., 2010, Escanilla et al., 2012).

#### Experimental procedure

Behavioral testing took place in a modified mouse cage with a start and a testing chamber separated by an opaque removable plexi glass door. Mice were put into the start chamber with the door closed and petri dishes with sand-odor mix were placed into the test chamber with a sugar pellet in the rewarded odor dish. The divider was openend and mice were allowed to dig in the dishes to retrieve the sugar reward. The time delay to dig in the correct dish was recorded by hand and later double checked on the video trace. Mice were shaped to dig using odorset 1 (Supplementary Table 1) until they consistently retrieved the reward for 18 out of 20 trials. For long term memory tests, each mouse was trained for 20 trials using two scented dishes, one rewarded one not, and tested on 5 trials 24 hours later. Odorsets 1&2 were used for shaping and odorsets 3-6 were used for the experimental trials. Each mouse was tested on each drug condition (saline, alpha1 blocker, alpha2 blocker, beta blocker, all blockers) with a different odorset; the order of drug conditions was pseudo randomized and counterbalanced among mice. For specificity testing, each mouse was trained on a odor-reward association with a 2-carbon odor (C) for 4, 8 or 12 trials paired with an unscented dish, immediately followed by unrewarded test trials with the conditioned odor (C), two similar odors (C+1&C+2) and one unrelated odor (X) (Supplementary Table 1). Each mouse was trained and tested under each drug condition and number of training trials with a different odorset, with order of drug conditions, number of trials and odor sets randomized and counterbalanced.

#### Data analysis

To test for the role of NE receptors in acquisition and 24 hour recall, we used repeated measure ANOVAs with latency to dig in the correct dish as dependent variable, drug group as between subjects factor and trial block or trial number as within subjects factor (alpha = 0.05), followed by pairwise comparisons between trial blocks to assess acquisition and recall using Wilk’s Lambda. For specificity testing, data were analyzed using a repeated measures analysis with drug group (saline or phent) and number of training trials (4, 8 or 12) as between subjects factors and digging times in response to unrewarded test odors (C, C1, C2 and X) as within subjects factor, followed by pairwise comparisons between digging responses in the conditioned and novel odors for each group (Wilk’s Lambda). All data analysis was performed in SPSS.

### Computational modeling

Computational modeling of the olfactory bulb followed the outline presented in Linster and Kelsch 2019, with detailed equations and parameter sets described in Supplementary methods and the associated Supplementary Table 2. The modeled OB network incorporates five neuron types: olfactory sensory neurons (OSNs), MCs, external tufted cells (ETs), periglomerular cells (PGs), and granule cells (GCs). Each group is composed of 100 neurons organized in functional columns with connectivity parameters specified in Supplementary Table 2. MCs make synapses with 25% of GCs (*p*_MC-GC_ = 0.25) and GCs make inhibitory local synapses onto MCs only. NE modulation to the OB was modeled according to the principles we discovered previously in brain slice and computational experiments (Nai et al., 2009, Nai et al., 2010, Linster et al., 2011); here, we simulated a high dosage of NE resulting in a dominance of alpha1 receptor effects on GC and MCs, which is also in agreement with results from the pharmacological experiments presented here. Briefly, NE alpha1 modulation increases MC excitability with no change in membrane voltage or spontaneous activity, and increases GC activation with an increase in voltage and spontaneous activity (Nai et al. 2009; 2010; Linster et al. 2011). Learning an odor reward association was modeled by projecting MC outputs in response to a conditioned odor (CS) to a response neuron (RN) which also received “reward” information (unconditioned stimulus (US), Figure 3A). Excitatory synapses between MCs and the RN underwent activity dependent synaptic plasticity when reward was present as well as a slow exponential decay in synaptic strength when reward was not present. This exponential decay had a time constant of 10 days, which resulted in memory durations similar to those observed experimentally. Reward association learning was simulated as follows. For each simulated “trial block” (5 trials of 30 seconds each), the conditioned odor was paired with reward (activation of RN by US) which resulted in changes in synaptic weights between MCs and RN. After each trial block we then set reward to zero and presented the conditioned odor C, two overlapping odors with varying degree of overlap (C+1, C+2; 78% resp. 34% correlation with C) simulating the odors used in Experiment 3 and an unrelated odor (X; −0.42 correlation with C) for one simulated trial and computed the resulting activation of the RN (Devore et al., 2014). This was repeated over the course of 4 trial blocks (20 trials total) to test to what degree the specificity of the association evolved as a function of learning and how this dependent on the presence of NE. Memory duration in the model was assessed by presenting the conditioned odor at intervals of 1 hour simulated time interleafed by 1 hour “forgetting” (see supplementary methods) in the model. We ran 10 different instances of the model, each initialized with a different seed for the random number generator. All parameters in the model are chosen from a uniform distribution +/− 10% around the mean value specified in Supplementary Table 2. To assess memory duration, we statistically compared MN response amplitudes (spiking probabilities) during pre-acquisition testing to that during each segment of the 24 hour forgetting period. To assess memory specificity, we statistically compared RN response magnitudes to conditioned and test odors after each trial block during acquisition.

---

**Supplementary Table 1.** Odors and dilutions used for behavioral experiments. All odorants were diluted in mineral oil to an approximate vapor partial pressure of 10 Pa. 60 uL of odorant was pipetted into the bedding used during behavioral training and testing before each trial.

| Odorsets | Odorant 1<br>(% vol/vol) | Odorant 2<br>(% vol/vol) |
| --- | --- | --- |
| 1 | Decanal<br>(17.76) | +Limonene<br>(2.0) |
| 2 | Anisole<br>(0.5) | Amyl Acetate<br>(0,7) |
| 3 | Pentanal<br>(0.07) | Butanal<br>(0,02) |
| 4 | Hexanoic acid<br>(14.9) | Pentanoic acid<br>(4.5) |
| 5 | Heptanol<br>(8.4) | Octanol<br>(26.7) |
| 6 | Butyl<br>Pentanoate<br>(5.7) | Butyl<br>Hexanoate<br>(16.3) |
| 7 | Butanol<br>(0.2) | Pentanol<br>(0.7) |

***Odorants and dilutions used for specificity testing***
| Odorsets | C<br>(% vol/vol) | C+1<br>(% vol/vol) | C+2<br>(% vol/vol) | X<br>(% vol/vol) |
| --- | --- | --- | --- | --- |
| 8 | Propanoic<br>acid<br>(0.33) | Butanoic acid<br>(1.3%) | Pentanoic<br>acid<br>(4.5) | 3-heptanone<br>(0.6) |
| 9 | Hexyl<br>acetate<br>(2.28) | Amyl acetate<br>(0.7) | Butyl acetate<br>(0.2) | Anisole<br>(0.5) |
| 10 | Pentanol<br>(0.74) | Hexanol<br>(2.6) | Heptanol<br>(8.4) | Benzylamine<br>(2.9) |
| 11 | Hexanoic<br>acid<br>(14.9) | Heptanoic acid<br>(46.8) | Octanoic<br>acid<br>(13.7) | Neryl acetate<br>(16.4) |
| 12 | Pentyl<br>butyrate<br>(5.8) | Hexyl butyrate<br>(16.3) | Heptyl<br>butyrate<br>(46.0) | +Carvone<br>(47.2) |
| 13 | Hexanal<br>(0.24) | Heptanal<br>(0.72) | Octanal<br>(1.48) | Trans-2-hexenyl<br>acetate<br>(26.7) |

**Supplementary Table 2.** Computational modeling parameters. Membrane time constant: τ resting membrane potential: V_rest_; spiking threshold: θ^min^; saturation threshold: θ^max^; synaptic weight: w; reversal potential : E_N_; rise time : τ_1_; decay time : τ_2_; after-hyperpolarization magnitude : A^ahc^; calcium accumulation time constant : τ^ahc^.

|  |  |
| --- | --- |
| Olfactory Sensory Neuron (OSN) | $\tau = 1\text{ms}$; $V_{rest} = -65\text{ mV}$; $\theta^{min} = -65\text{mV}$; $\theta^{max} = -55\text{mV}$. |
| Mitral | $\tau = 5\text{ms}$; $V_{rest} = -65\text{ mV}$; $\theta^{min} = -64\text{mV}$; $\theta^{max} = -57/61\text{mV}^*$ |
| Periglomerular (PG) | $\tau = 2\text{ms}$; $V_{rest} = -65\text{ mV}$; $\theta^{min} = -65\text{mV}$; $\theta^{max} = -60\text{mV}$ |
| Granule (Gr) | $\tau = 4\text{ms}$; $V_{rest} = -65\text{ mV}$ ; $\theta^{min} = -65.2/-66\text{mV}$; $\theta^{max} = -60\text{mV}$ |
| External tufted (ET) | $\tau = 2\text{ms}$; $V_{rest} = -65\text{ mV}$ ; $\theta^{min} = -65\text{mV}$; $\theta^{max} = -60\text{mV}$. |
| Pyramidal (Pyr) | $\tau = 10\text{ms}$; $V_{rest} = -65\text{ mV}$; $\theta^{min} = -62\text{mV}$; $\theta^{max} = -55\text{mV}/-60\text{mV}^*$; |
| OSN to PG | $w = 0.003$; $E_N = +70\text{mV}$; $\tau_1 = 1\text{ms}$; $\tau_2 = 2\text{ms}$ |
| OSN to Mi (apical) | $w = 0.006$; $E_N = +70\text{mV}$; $\tau_1 = 1\text{ms}$; $\tau_2 = 2\text{ms}$ |
| OSN to ET(apical) | $w = 0.0006$; $E_N = +70\text{mV}$; $\tau_1 = 1\text{ms}$; $\tau_2 = 2\text{ms}$ |
| PG to Mi (apical) | $w = 0.003$; $E_N = -5\text{mV}$; $\tau_1 = 2\text{ms}$; $\tau_2 = 4\text{ms}$ |
| ET to Mi (apical) | $w = 0.0015$; $E_N = 70\text{mV}$; $\tau_1 = 1\text{ms}$; $\tau_2 = 2\text{ms}$ |
| Mi (soma) to Gr | $w = 0.004$ $E_N = +70\text{mV}$; $\tau_1 = 1\text{ms}$; $\tau_2 = 2\text{ms}$ ; $p = 0.2$; |
| Gr to Mi (soma) | $w = 0.004$; $E_N = -5\text{mV}$; $\tau_1 = 2\text{ms}$; $\tau_2 = 4\text{ms}$ ; local only |
| Mi (soma) to RN | $w_{naive} = 0.001$; $E_N = +70\text{mV}$; $\tau_1 = 1\text{ms}$; $\tau_2 = 2\text{ms}$; |

\* different values are without/with NE modulation, respectively;

**Supplementary Figure 1.**
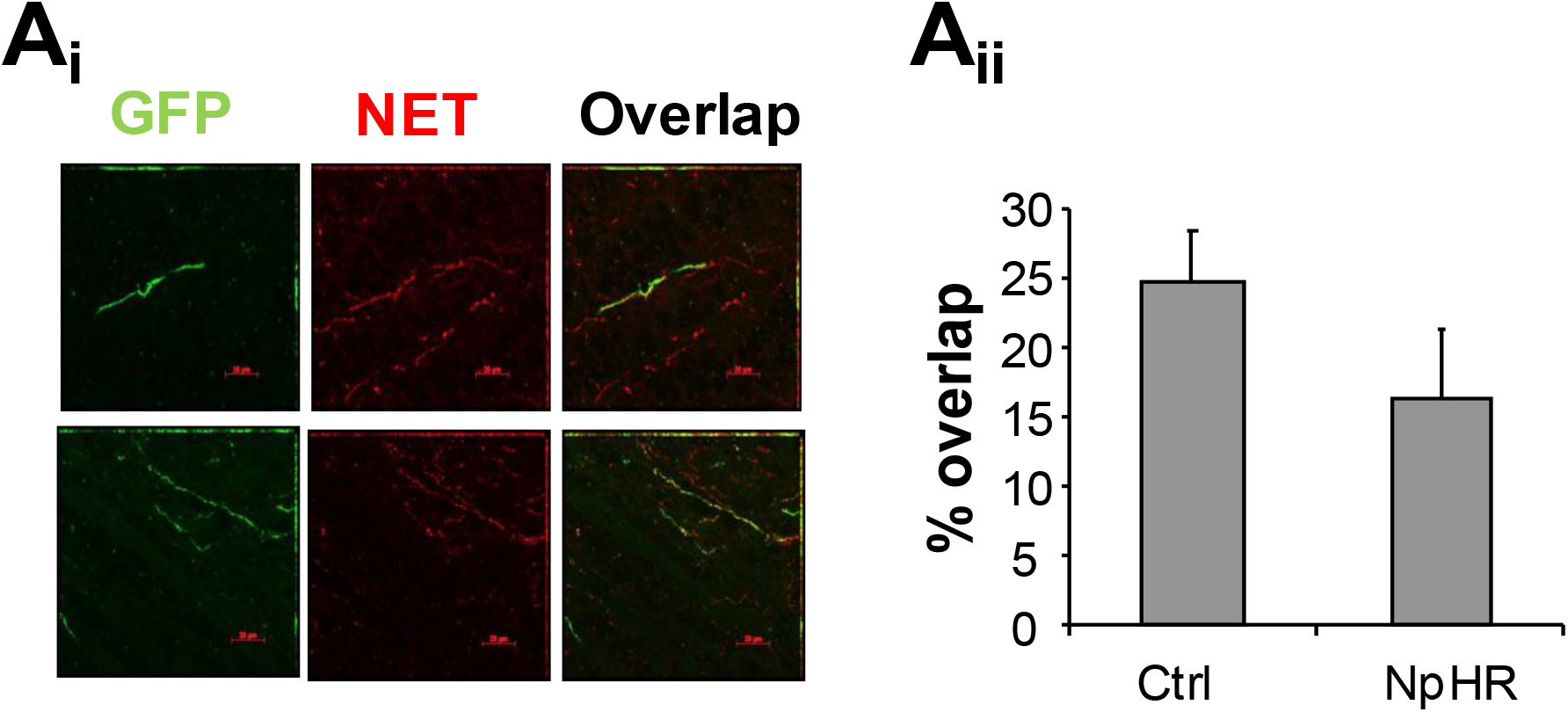
Viral transfection in NE fibers terminating in the OB. **Ai.** GFP, NET and overlap between GFP and NET images in the OB. **Aii.** The amount of overlap between GFP and NET expressing fibers was similar in Ctrl (n=5) and NpHR (n=7) mice (F(1, 10) = 1.827; p = 0.206).

**Supplementary Figure 2.**
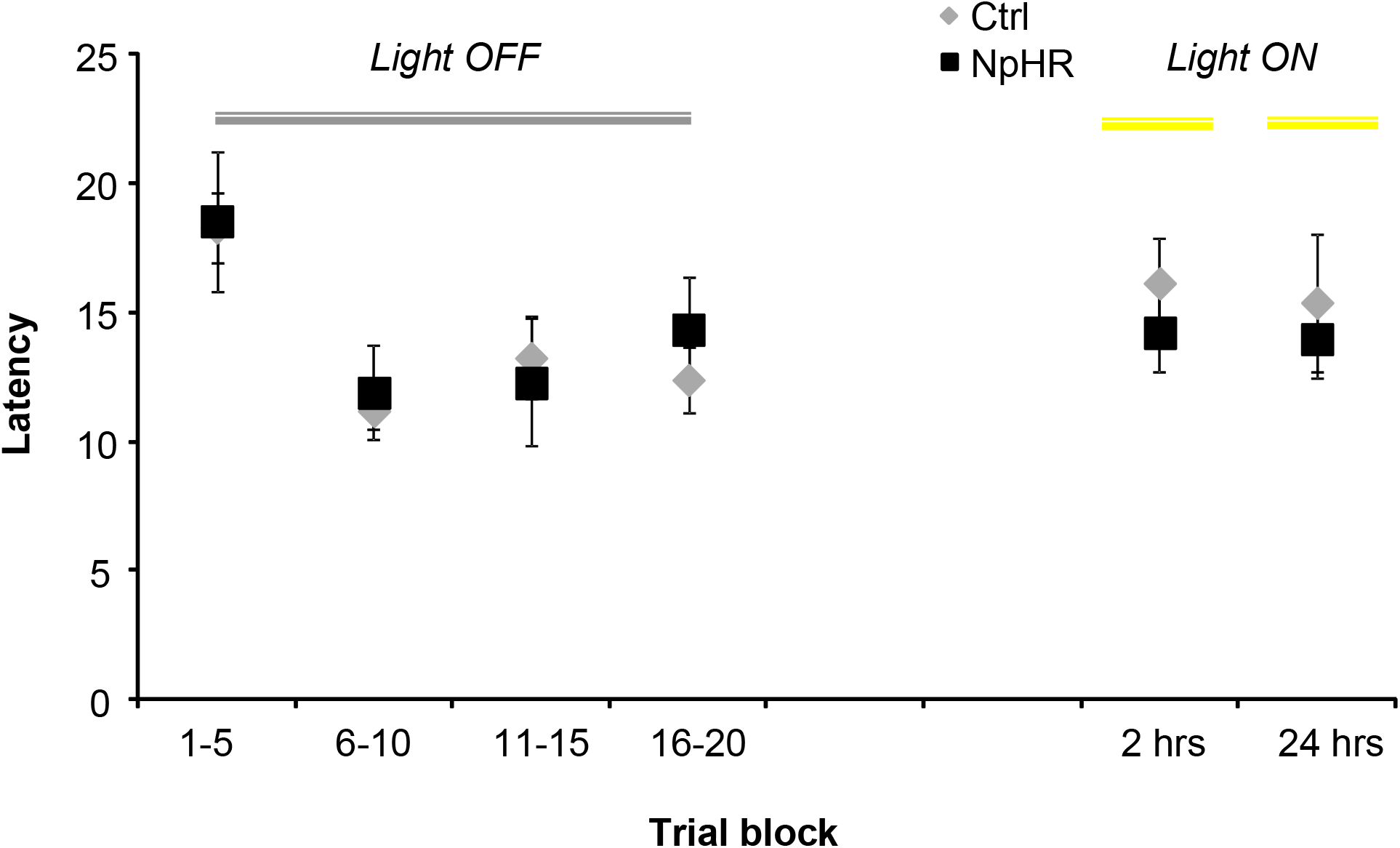
Recall control experiment. Mice were first trained for 20 trials without light stimulation (LIGHT OFF; gray line) and underwent recall trials at 2 hours and 24 hours with light stimulation (LIGHT ON; yellow line). We found a significant effect of trial block (F_trialblock_ (3, 7) = 6.000; P = 0.017) but no interaction with group (F_trialblock,group_ (3, 7) = 0.824; p=0.696) during acquisition showing that while both groups learned there was no difference in their acquisition curves. In contrast to findings from the original experiment, with respect to long-term memory, there was no effect of trial block (short-term versus long-term; F(1, 9) = 0.938; p = 0.358) or interaction with experimental group (F(1, 9) = 0.010; p = 0.922) indicating that both groups perform similarly in both recall situations. Neither group displayed significantly longer delays at 24 hours than during the last acquisition trials (respectively, p = 0.669 and p = 0.759).

**Supplementary Figure 3.**
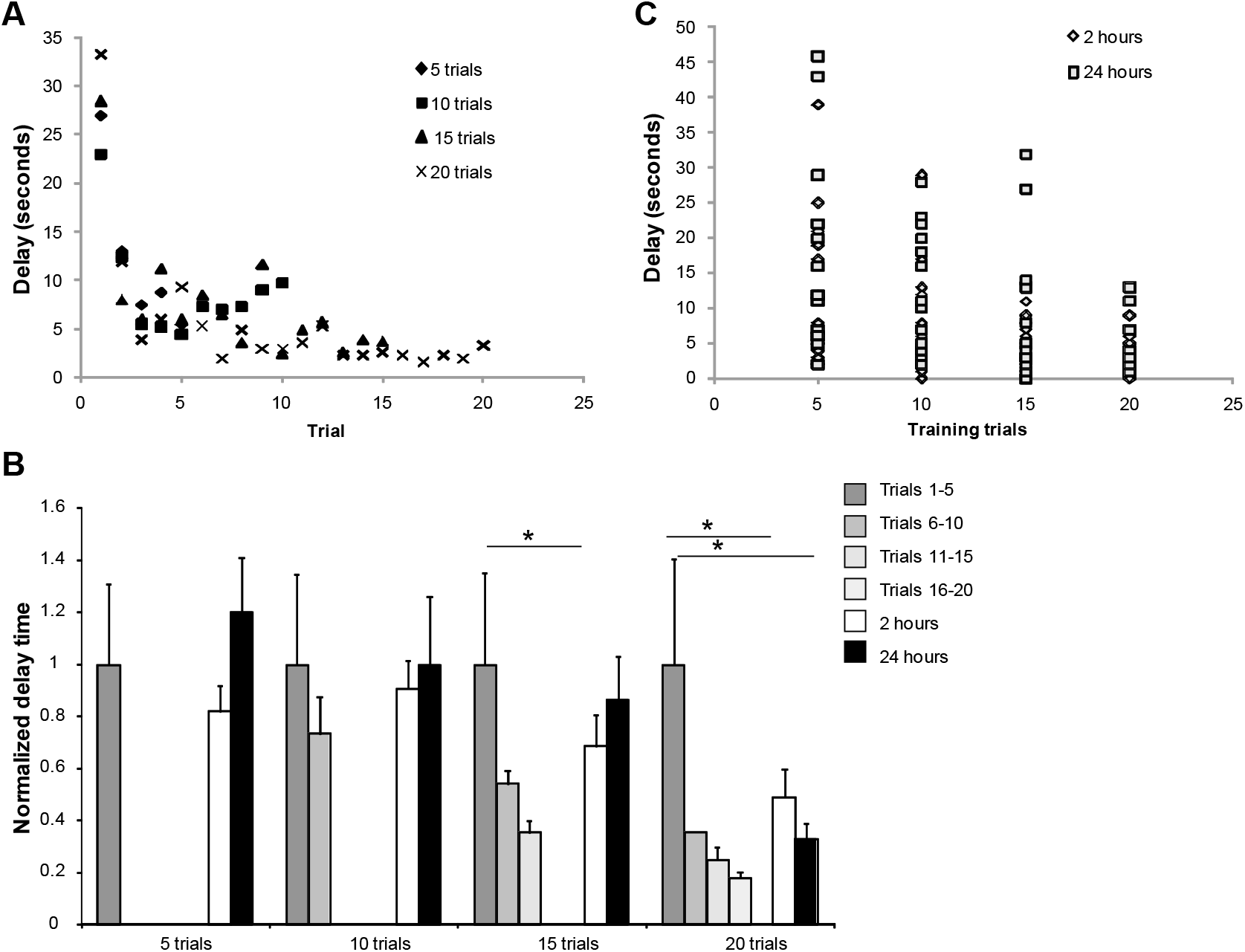
Memory stability, expressed by duration, increases with amount of training. In a separate experiment we tested the idea that amount of training would increase stability also with respect to memory duration. Towards this goal, mice were trained in the simultaneous go – no go task to retrieve a sugar reward from one odor in an easily discriminated pair and the delay to find the correct dish recorded as a measure for learning (odorsets 8,9,10&11 from Table 1 were used). Mice were trained for 5, 10, 15 or 20 trials (with different odorsets) and tested 2 hours or 24 hours later. In each case, mice learned during the acquisition trials as evidenced by decreasing delays to find the correct dish (significant negative correlation between trial number and delay to dig time (R=-0.611; p < 0.001 with Pearson’s R; (**A**). Memory for the learned odor is indicated when mice need significantly less time to find the rewarded odor during the 5 test trials than during the initial five acquisition trials. A repeated measures analysis with delay times during the initial five trials and trials after 2 and 24 hours showed a significant effect of trial set after 15 and 20 training trials only (F(2, 14) = 6.174; p = 0.012 and F(2, 11) = 4.354; p = 0.04). Mice remembered the odor after 2 hours after 15 and 20 trial training (significant decrease of delay at 2 hour test (p < 0.05) and after 24 hours with 20 training trials only (p = 0.007; **B**). These data show a strong effect of amount of training on long term memory of an odor reward association, also indicated by a strong negative correlation between the number of training trials and the delay to find the correct odor during the 2 hours and 24 hour tests (R=-0.332; p = 0.004 and R=-0.353; p = 0.002; (**C**)). Together with results from our previous experiments, these data further support the idea that more training leads to stronger memory and bulbar NE can compensate for training or “speed up” learning.

**Supplementary Figure 4.**
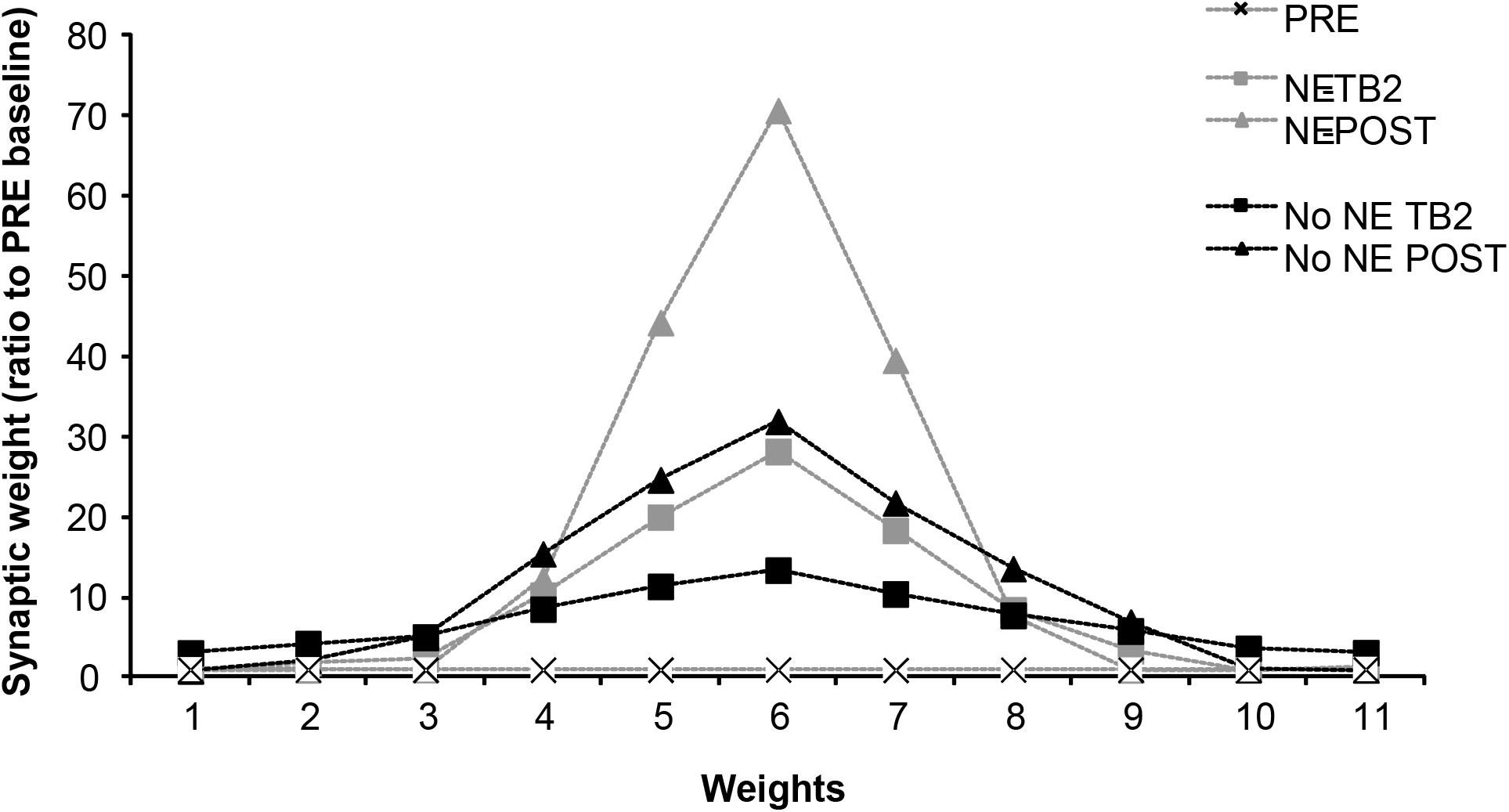
Synaptic weight changes as a function of learning and NE inputs in the model. The graph shows synaptic weights normalized to the initial weights and ordered by amplitude: pre-conditioning (PRE), after two trial blocks (TB2) and post conditioning (after 4 trial blocks) with NE and without NE (no NE). As weights grow with learning, the distribution becomes more narrow and specific to the conditioned odor due to competition between synapses in the learning rule. Weight distributions are more narrow when acquisition is done in the presence of NE.

## Supplementary Methods : Computational modeling

### Network architecture

The modeled OB network incorporates five neuron types: olfactory sensory neurons (OSN), mitral cells (MC), external tufted cells (ET), periglomerular cells (PG) and granule cells (GC). Each group is composed of 100 neurons organized in functional columns. MCs make synapses with 25% of GCs (p_MC-GC_=0.25) and GCs make inhibitory local synapses only (see (McIntyre and Cleland, 2016). To assess associations between odors (CS) and reward (US) a response neuron (RN) was added which received excitatory synaptic inputs with very low initial weights from all MCs and underwent activity dependent synaptic plasticity when US and CS were present at the same time. During post – acquisition, when no US was present these synapses underwent a slow exponential decay back to baseline values. Response magnitude of the RN were measured as instantaneous spiking probabilities.

### Neurons and synapses

Our model is composed of single compartment leaky integrate-and-fire neurons, with the exception of MC which are modeled as two compartments. Changes in membrane voltage v(t) over time in each compartment are described by eq. 1:

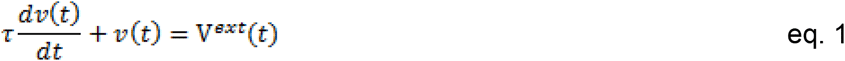

where τ is the membrane time constant and *V^ext^*(*t*) is the voltage change resulting from external inputs (synaptic or sensory).

Each one of the voltage changes due to external inputs *V^ext^* is a result of the synaptic strength of the connection from neuron *j* to neuron *i* (*w_ij_*) and the respective synaptic conductance in cell i at time t (*g_i_*(*t*)). *E_N,ij_* is the Nernst potential of the synaptic current and *v_i_*(*t*) is the membrane potential of the postsynaptic neuron *i*, as described in eq. 2:

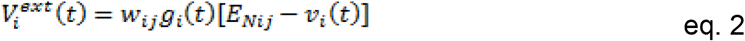

The communication between neurons happens via discrete spikes. The spiking output *F(v)* of a given neuron *i* is a function of its membrane potential *v* and the minimal threshold and saturation threshold of the output function, *θ^min^* and *θ^max^*. Where *F_i_*(*v*) = 0 if *v*≤*θ^min^* and *F_i_*(*v*) = 1 if *v*≥*θ^max^* and Fi(v) increase linearly between θ_min_ and θ_max_

*F*_i_(*v*) defines their instantaneous firing probability and OXT modulation decreases θ_max_ to increase excitability. The time course of the conductance change is calculated as: eq.4

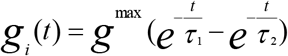

where *g^max^* is a constant with no unit representing the maximum conductance of a given channel and is equal to 1 (synaptic strength is scaled by the synaptic weight w), while τ_1_ and τ_2_ are the rising and falling times of this conductance. After firing, the spike of each spiking-neuron is reset to V_rest_.

In the simulations presented here, simulated exposure to an odorant induced activity dependent plasticity of synapses from MC to the RN. Synaptic strengths were first calculated from the parameters given in Supplementary Table2. During simulated trial blocks, synapses between MCs and the RN underwent synaptic potentiation:

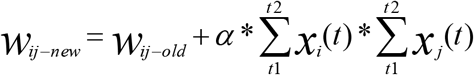

where *w_ij_* is the synaptic strength between the presynaptic MC and postsynaptic RN,α(α = 0.01) is the rate of potentiation and *x_j_* and *x_i_* are the total numbers of spikes emitted by the pre and postsynaptic cells during the preceding sniff cycles between t1 and t2. The synaptic weights also undergo postsynaptic normalization after the weight change has been computed, with the sum of synaptic weights from MC to RN staying constant (the sum was determined by the sum of initial weights); this creates competition between synaptic weights over the course of training and leads to increasing specificity of synaptic weights to the odor used for conditioning. Post acquisition synaptic weights decay exponentially with a long time constant (10 hours) to simulate forgetting in the absence of US-CS pairings. Any synaptic weight below 0.0 is reset to a value within the initial low random distribution. This time constant was adjusted to result in a long term memory of at least 24 hours for simulations in which NE was present during acquisition.

### Implementation

All simulations were implemented using the C programming language in a Linux environment (Ubuntu 14.04 LTS x64) on an Intel desktop computer, with Euler integration method for the differential equations with a time step of 1ms. Parameters were chosen from uniform random distributions +/− 10% around the mean values given in Supplementary Table 2.

### Code Accessibility

The code/software described in the paper is freely available online at [add data base here]. The code is available as Extended Data (Extended data 1).

## References

Chaudhury D, Escanilla O, Linster C (2009) Bulbar acetylcholine enhances neural and perceptual odor discrimination. J Neurosci 29:52–60.

Cleland TA, Morse A, Yue EL, Linster C (2002) Behavioral models of odor similarity. Behav Neurosci 116:222–231.

Cleland TA, Narla VA (2003) Intensity modulation of olfactory acuity. Behav Neurosci 117:1434–1440.

Cleland TA, Narla VA, Boudadi K (2009) Multiple learning parameters differentially regulate olfactory generalization. Behav Neurosci 123:26–35.

de Almeida L, Reiner SJ, Ennis M, Linster C (2015) Computational modeling suggests distinct, location-specific function of norepinephrine in olfactory bulb and piriform cortex. Front Comput Neurosci 9:73.

Devore S, de Almeida L, Linster C (2014) Distinct roles of bulbar muscarinic and nicotinic receptors in olfactory discrimination learning. J Neurosci 34:11244–11260.

Devore S, Linster C (2012) Noradrenergic and cholinergic modulation of olfactory bulb sensory processing. Front Behav Neurosci 6:52.

Doucette W, Milder J, Restrepo D (2007) Adrenergic modulation of olfactory bulb circuitry affects odor discrimination. Learn Mem 14:539–547.

Escanilla O, Alperin S, Youssef M, Ennis M, Linster C (2012) Noradrenergic but not cholinergic modulation of olfactory bulb during processing of near threshold concentration stimuli. Behav Neurosci 126:720–728.

Escanilla O, Arrellanos A, Karnow A, Ennis M, Linster C (2010) Noradrenergic modulation of behavioral odor detection and discrimination thresholds in the olfactory bulb. Eur J Neurosci 32:458–468.

Escanilla O, Mandairon N, Linster C (2008) Odor-reward learning and enrichment have similar effects on odor perception. Physiol Behav 94:621–626.

Gentry RN, Schuweiler DR, Roesch MR (2019) Dopamine signals related to appetitive and aversive events in paradigms that manipulate reward and avoidability. Brain Res 1713:80–90.

Gradinaru V, Zhang F, Ramakrishnan C, Mattis J, Prakash R, Diester I, Goshen I, Thompson KR, Deisseroth K (2010) Molecular and cellular approaches for diversifying and extending optogenetics. Cell 141:154–165.

Guerin D, Peace ST, Didier A, Linster C, Cleland TA (2008) Noradrenergic neuromodulation in the olfactory bulb modulates odor habituation and spontaneous discrimination. Behav Neurosci 122:816–826.

Kermen F, Chakirian A, Sezille C, Joussain P, Le Goff G, Ziessel A, Chastrette M, Mandairon N, Didier A, Rouby C, Bensafi M (2011) Molecular complexity determines the number of olfactory notes and the pleasantness of smells. Sci Rep 1:206.

Kermen F, Midroit M, Kuczewski N, Forest J, Thevenet M, Sacquet J, Benetollo C, Richard M, Didier A, Mandairon N (2016) Topographical representation of odor hedonics in the olfactory bulb. Nat Neurosci 19:876–878.

Linster C (2019) Cellular and network processes of noradrenergic modulation in the olfactory system. Brain Res 1709:28–32.

Linster C, Escanilla O (2019) Noradrenergic effects on olfactory perception and learning. Brain Res 1709:33–38.

Linster C, Hasselmo ME (1999) Behavioral responses to aliphatic aldehydes can be predicted from known electrophysiological responses of mitral cells in the olfactory bulb. Physiol Behav 66:497–502.

Linster C, Nai Q, Ennis M (2011) Nonlinear effects of noradrenergic modulation of olfactory bulb function in adult rodents. J Neurophysiol 105:1432–1443.

Mandairon N, Ferretti CJ, Stack CM, Rubin DB, Cleland TA, Linster C (2006) Cholinergic modulation in the olfactory bulb influences spontaneous olfactory discrimination in adult rats. Eur J Neurosci 24:3234–3244.

Mandairon N, Kuczewski N, Kermen F, Forest J, Midroit M, Richard M, Thevenet M, Sacquet J, Linster C, Didier A (2018) Opposite regulation of inhibition by adult-born granule cells during implicit versus explicit olfactory learning. Elife 7.

Mandairon N, Peace S, Karnow A, Kim J, Ennis M, Linster C (2008) Noradrenergic modulation in the olfactory bulb influences spontaneous and reward-motivated discrimination, but not the formation of habituation memory. Eur J Neurosci 27:1210–1219.

Mandairon N, Sultan S, Rey N, Kermen F, Moreno M, Busto G, Farget V, Messaoudi B, Thevenet M, Didier A (2009) A computer-assisted odorized hole-board for testing olfactory perception in mice. J Neurosci Methods 180:296–303.

Manella LC, Alperin S, Linster C (2013) Stressors impair odor recognition memory via an olfactory bulb-dependent noradrenergic mechanism. Front Integr Neurosci 7:97.

Manella LC, Petersen N, Linster C (2017) Stimulation of the Locus Ceruleus Modulates Signal-to-Noise Ratio in the Olfactory Bulb. J Neurosci 37:11605–11615.

McBurney-Lin J, Lu J, Zuo Y, Yang H (2019) Locus coeruleus-norepinephrine modulation of sensory processing and perception: A focused review. Neurosci Biobehav Rev 105:190–199.

McIntyre AB, Cleland TA (2016) Biophysical constraints on lateral inhibition in the olfactory bulb. J Neurophysiol 115:2937–2949.

Moreno MM, Bath K, Kuczewski N, Sacquet J, Didier A, Mandairon N (2012) Action of the noradrenergic system on adult-born cells is required for olfactory learning in mice. J Neurosci 32:3748–3758.

Nai Q, Dong HW, Hayar A, Linster C, Ennis M (2009) Noradrenergic regulation of GABAergic inhibition of main olfactory bulb mitral cells varies as a function of concentration and receptor subtype. J Neurophysiol 101:2472–2484.

Nai Q, Dong HW, Linster C, Ennis M (2010) Activation of alpha1 and alpha2 noradrenergic receptors exert opposing effects on excitability of main olfactory bulb granule cells. Neuroscience 169:882–892.

Parikh V, Sarter M (2008) Cholinergic mediation of attention: contributions of phasic and tonic increases in prefrontal cholinergic activity. Ann N Y Acad Sci 1129:225–235.

Sarter M, Gehring WJ, Kozak R (2006) More attention must be paid: the neurobiology of attentional effort. Brain Res Rev 51:145–160.

Tong MT, Kim TP, Cleland TA (2018) Kinase activity in the olfactory bulb is required for odor memory consolidation. Learn Mem 25:198–205.

